# Assessing the translation of AI-prioritized genome-derived peptide fragments into validated antimicrobial candidates

**DOI:** 10.64898/2026.08.25.747168

**Authors:** Sebastian Ojeda, Paula Avila, Stiven Castellanos, Paloma Lemaitre, Valeria Ruiz-Ramírez, Marcela Manrique-Moreno, Adriana Marcela Celis Ramírez, Pablo Arbeláez, Chad Leidy, Carolina Muñoz-Camargo

## Abstract

The emergence of antibiotic-resistant pathogens such as *Staphylococcus aureus* demands accelerated antimicrobial discovery strategies. Artificial intelligence (AI) enables large-scale inference of candidate antimicrobial peptides (AMPs), yet experimental validation remains essential to determine whether predictions translate into biological function. Genome-guided mining, rather than unconstrained or randomly generated sequence exploration, offers a biologically grounded search space derived from organisms shaped by ecological and evolutionary pressures. Here, we evaluate this principle using *Malassezia furfur*, a skin-associated yeast that coexists with bacterial colonizers such as *S. aureus*, as a genomic source for AI-prioritized antimicrobial candidates. Candidate fragments were generated from two *M. furfur* genomes, filtered by physicochemical properties, prioritized with deep-learning AMP predictors, synthesized, and experimentally characterized. Selected peptides underwent cross-kingdom antimicrobial screening against *S. aureus*, combining kinetic growth and ultrastructural assays, complemented by *in silico* structural prediction, lipid-membrane interaction analysis, and human keratinocyte cytotoxicity evaluation. AI-guided genomic mining enriched biologically motivated sequence space for peptides with measurable antimicrobial activity, while revealing biases and generalizability limits of AI-based AMP inference. Closing the loop between genome-derived candidate generation, AI-based inference, synthesis, and functional characterization, this study provides an experimental assessment of model-guided AMP discovery and a reproducible route from computational prediction to validated antimicrobial candidates.

## Introduction

Antibiotics are foundational to modern medicine, enabling routine surgical procedures, immunosuppressive therapies, and the treatment of common infections. However, widespread misuse and overuse have accelerated the emergence of antimicrobial resistance (AMR), allowing pathogens to endure drug exposure and compromising clinical outcomes across infectious diseases. Recent global health projections estimate that antimicrobial-resistant infections could cause tens of millions of deaths in the coming decades, with disproportionate impact across several regions worldwide^1^. Beyond morbidity and mortality, AMR also imposes a substantial economic burden by increasing healthcare costs and threatening productivity at the population scale^2^. This escalating crisis has intensified the search for alternative anti-infective modalities capable of controlling pathogens while reducing the likelihood of rapid resistance development.

Among the most promising alternative modalities are antimicrobial peptides (AMPs), naturally occurring or synthetically designed short molecules typically composed of 5–50 amino acids with broad antimicrobial properties. Unlike many traditional antibiotics that act on discrete intracellular targets, AMPs frequently exert activity through mechanisms harder for pathogens to evade, often involving membrane interaction, permeability changes, or multi-target disruption, thereby lowering the probability of resistance emerging from single-point mutations^3^. Nonetheless, translating AMP potential into clinically useful candidates remains difficult: discovery is constrained by the enormous combinatorial sequence space, while development is challenged by instability in physiological environments, variable potency across assay conditions, and safety concerns such as cytotoxicity^4^. Consequently, AMP research increasingly requires methodologies capable of prioritizing candidates at scale while connecting predictions to rigorous experimental validation.

Artificial intelligence (AI) has emerged as a key accelerator of AMP discovery, enabling high-throughput screening and prioritization across large peptide repertoires^5^. Modern deep-learning predictors learn sequence and feature-level patterns associated with antimicrobial activity from curated databases, such as the Antimicrobial Peptide Database (APD)^6^, the Collection of Antimicrobial Peptides (CAMP)^7^, and others, offering scalable priors difficult to encode as hand-designed rules^8^. Nevertheless, AI-driven discovery faces persistent translational challenges. Training datasets are inherently skewed, as they overrepresent specific target organisms and clinical assay conditions. For instance, within the Antimicrobial Peptide Database (APD6), a large majority of cataloged entries (83.56%) target bacterial pathogens, while a much smaller fraction (27.66%) possess documented antifungal activity^6^; these categories are not mutually exclusive, but the disparity is nonetheless illustrative of a bacterial-heavy skew in curated AMP resources. This severe taxonomic asymmetry is further compounded by synthesis biases; major repositories such as the Database of Antimicrobial Activity and Structure of Peptides (DBAASP), which currently catalogs 25,069 entries, reveal that the overwhelming majority of studied sequences are synthetic constructs rather than natural, translationally diverse ribosomal or non-ribosomal variants^9^. Consequently, predictive tools are frequently biased toward familiar, overrepresented chemical spaces and sequence motifs, drastically degrading their performance and reliability when translating inference onto out-of-distribution biological sources. Moreover, many studies emphasize cross-validation metrics without systematically testing whether high-confidence predictions retain activity when transferred to new biological sources or experimental settings. As a result, experimental validation remains a critical bottleneck: without it, it is difficult to determine whether model inference reflects generalizable biological signal or dataset-specific correlations. Genome-guided AMP discovery offers a principled route to mitigate this gap: by restricting the search to biologically plausible sequences within a given organism, genomic mining narrows the candidate space to molecules that may have been shaped by ecological pressures and host-associated selection^10^. Yet even within a single genome, naive enumeration yields an immense number of candidates, making principled triage essential; physicochemical heuristics provide an effective first pass^11^, but must be complemented by AI-based prioritization to reduce false positives and improve translational confidence.

The selection of *Malassezia furfur* as a genomic source is motivated by its ecological role within the human skin microbiome. *Malassezia* spp. are lipid-dependent yeasts that represent a major fungal component of the cutaneous microbiome and colonize sebaceous regions throughout the body^12^. In this environment, they coexist with bacterial commensals and opportunistic pathogens, including *Staphylococcus aureus*, a clinically relevant pathobiont associated with skin disorders such as atopic dermatitis and wound infections^13^. Their shared, resource-limited niche suggests that interspecies competition may involve secreted metabolites, lipases, or antimicrobial factors that influence microbial community structure and colonization resistance. This ecological context provides a plausible selective basis for exploring whether *M. furfur* encodes peptide fragments with antibacterial potential against *S. aureus*, making it a biologically motivated source for AMP discovery rather than an arbitrary genomic reservoir.

In this study, we test whether AI-guided genomic mining of *M. furfur* can identify peptide candidates with measurable antimicrobial activity and, more broadly, whether the peptides prioritized by the model demonstrate the predicted antimicrobial function in an experimental setting. We implement an integrated computational-to-experimental workflow in which candidate fragments are generated through *in silico* restriction-enzyme digestion of two *M. furfur* genomes, reduced through physicochemical filtering based on antimicrobial-like benchmarks—specifically length (10–50 amino acids), overall cationicity (net charge +3 to +7.5), and bounded amphipathic character (hydrophobic moment *≤* 0.6) (Figure 1a)—and prioritized using our previously developed and validated deep-learning predictor, AMPs-Net^14^ (Figure 1b). From this pipeline, based on their physicochemical properties, four candidates were prioritized for synthesis. We then determined their antibacterial activity against *S. aureus*, characterized the resulting ultrastructural morphology by electron microscopy, evaluated their antifungal activity against the source organism *M. furfur*, assessed lipid-membrane interactions and peptide secondary structure by FT-IR spectroscopy, and evaluated cytotoxicity in mammalian cell lines (Figure 1c).

**Figure 1.**
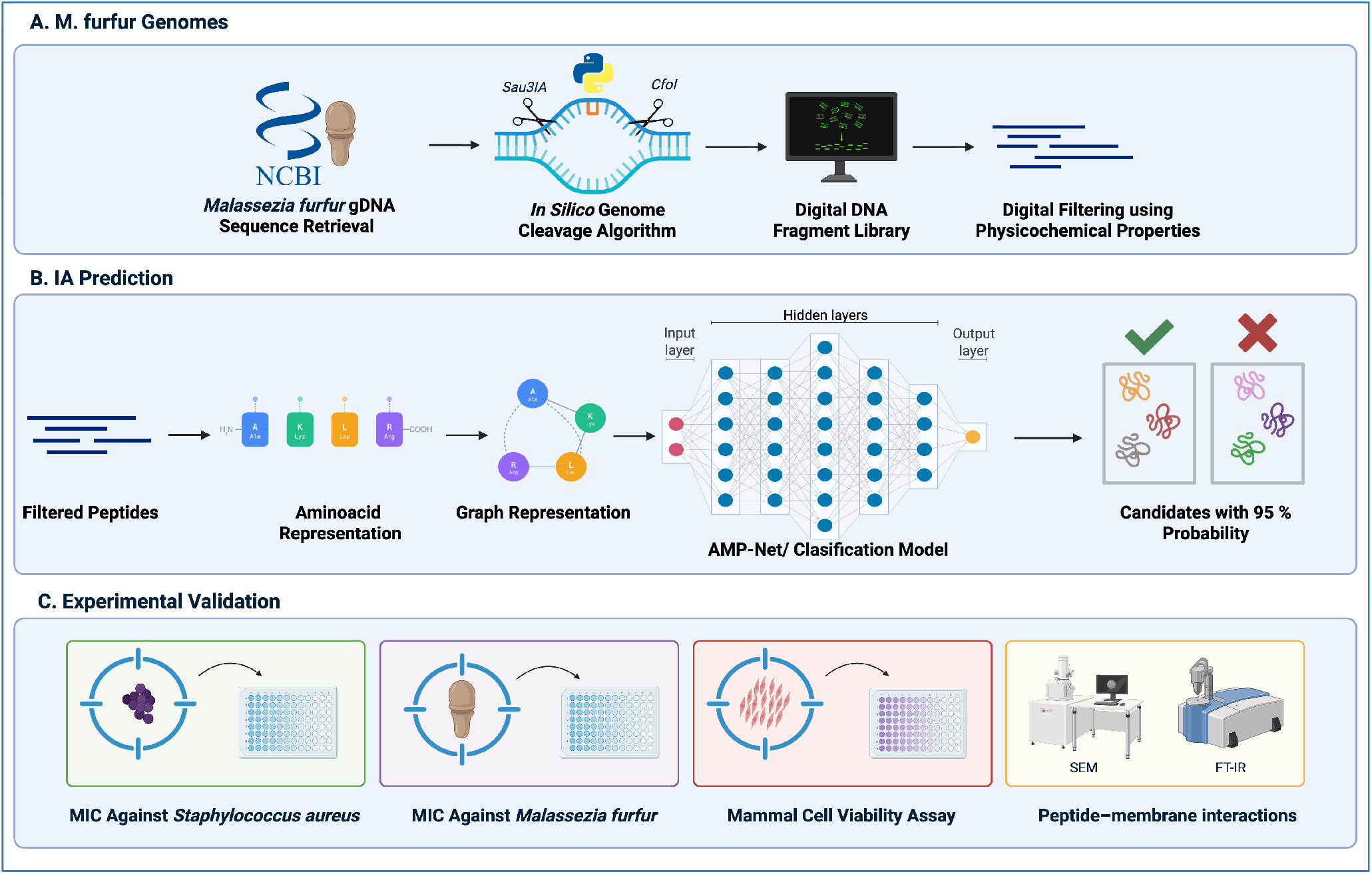
Overview of the AI-guided genome-mining and experimental validation pipeline. **(a)** *M. furfur* genomic sequences were retrieved, fragmented by *in silico* restriction-enzyme cleavage, and filtered using physicochemical criteria to generate a refined peptide library. **(b)** Filtered peptides were represented using sequence- and graph-based encodings and prioritized with the AMPs-Net classification model to select high-probability antimicrobial candidates. **(c)** Selected candidates were validated through antibacterial assays against *S. aureus*, activity testing against *M. furfur*, mammalian cell viability assays, scanning electron microscopy (SEM), and Fourier-transform infrared spectroscopy (FT-IR) analysis of peptide–membrane interactions.

## Results

### *M. furfur* Genome-Derived Peptide Discovery

We sought to determine whether the *M. furfur* genome contains peptide fragments with antibacterial potential against *S. aureus*. To this end, we implemented a genome-guided discovery workflow that integrates *in silico* genome fragmentation, AMP-inspired physicochemical triage, deep-learning prioritization (AMPs-Net), cross-strain conservation filtering, and final downselection for experimental feasibility^14,15^. This multi-stage strategy reduced an initially massive fragment space to a small set of candidates suitable for synthesis, structural analysis, and *in vitro* validation.

### *M. furfur* Genome Data Acquisition and *in silico* Fragmentation

We analyzed two *M. furfur* strains (CBS 1878 and 4DS) as independent genomic sources^16,17^. Using two strains serves as an internal robustness check because fragments that persist across both genomes are less likely to reflect strain-specific variation and more likely to represent conserved sequence determinants. To generate peptide candidates, both genomes were cleaved *in silico* using two restriction enzymes (*Sau3AI* and *CfoI*)^14,15,18^. This step yielded a large starting search space of 665,096 fragments across strains and enzymes. Fragment yield differed strongly by enzyme, with *CfoI* producing substantially more fragments (270,708 for CBS 1878 and 251,956 for 4DS) than *Sau3AI* (64,679 for CBS 1878 and 77,753 for 4DS) (Tables 1 and 2). These counts define the initial candidate space that must be narrowed to isolate fragments compatible with AMP-like properties.

**Table 1.**
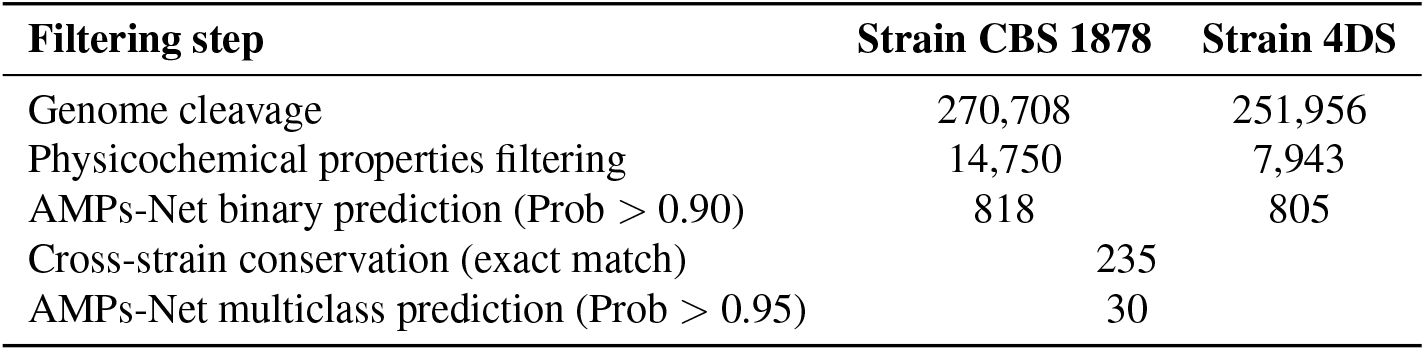
Filtering process for *M. furfur* strains CBS 1878 and 4DS using the restriction enzyme *CfoI*.

**Table 2.**
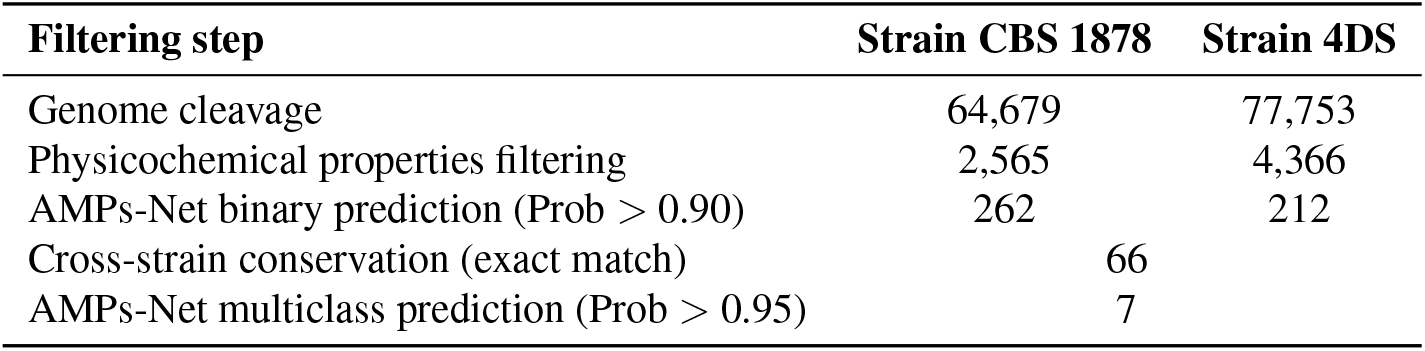
Filtering process for *M. furfur* strains CBS 1878 and 4DS using the restriction enzyme *Sau3AI*.

### Physicochemical Filtering of *M. furfur* Candidate Peptides

Because AMPs occupy a narrow region of physicochemical space, we next applied AMP-inspired constraints to remove fragments unlikely to be bioactive^11,19^. For each fragment we computed standard descriptors including length, net charge, hydrophobic moment, instability index, and isoelectric point^15^. We then retained fragments within an AMP-compatible length range (10–50 amino acids), exhibiting overall cationicity (net charge +3 to +7.5 at physiological pH), and bounded hydrophobic character (hydrophobic moment *≤* 0.6), reflecting the typical requirements for electrostatic attraction to the negatively charged bacterial membrane and the capacity to partition into lipid bilayers without compromising solubility or stability.

This stage produced a major contraction of the candidate space. For *CfoI*, physicochemical filtering reduced fragments from 270,708 to 14,750 (CBS 1878) and from 251,956 to 7,943 (4DS). For *Sau3AI*, it reduced fragments from 64,679 to 2,565 (CBS 1878) and from 77,753 to 4,366 (4DS) (Tables 1 and 2). Overall, fewer than 5% of the initial fragments satisfied these AMP-like constraints, highlighting that most genome-derived fragments are biophysically implausible as AMPs and underscoring the importance of triage prior to model-based prioritization.

### Deep-Learning–Based AMP Prediction with AMPs-Net

Fragments passing physicochemical triage were then prioritized using AMPs-Net, a deep-learning predictor trained to identify AMPs and to assign subclass probabilities (antifungal, antiviral, antiparasitic and antibacterial activity)^15^. Applying a stringent binary antimicrobial probability threshold (Prob *>* 0.90) retained 818 candidates from CBS 1878 and 805 from 4DS for the *CfoI*-derived set, and 262 (CBS 1878) and 212 (4DS) candidates for the *Sau3AI*-derived set (Tables 1 and 2). This step substantially reduced the search space while preserving candidates most likely to express AMP-like functional signatures according to the model.

To emphasize robust genomic signals and reduce the likelihood of strain-specific artifacts, we next retained only fragments present as an identical sequence in both strains. This conservation filter yielded 235 candidates for *CfoI* and 66 candidates for *Sau3AI* (Tables 1 and 2). Finally, we used the multiclass antibacterial output of AMPs-Net with a high-confidence threshold (Prob *>* 0.95), producing 30 antibacterial candidates for *CfoI* and 7 for *Sau3AI*. In total, the computational pipeline yielded 37 high-confidence antibacterial candidates encoded within the *M. furfur* genome.

### Selection of *M. furfur* -derived Candidate Peptides for Synthesis

While the 37 candidates represent a manageable computational output, experimental synthesis and testing of all peptides would be impractical. A final prioritization was therefore performed based on experimental feasibility and predicted structural stability, guided by a subset of physicochemical descriptors including net charge, hydrophobic moment, and predicted stability^20^. This refinement reduced the candidate set to seven peptides: PIAM11-4, PIAM18-5, PIAM21-4, PIAM21-7, PIAM23-3, PIAM23-4 and PIAM29-7 (PIAM: peptide identified artificially from *Malassezia*, where the first number denotes the number of amino acids and the number following the hyphen denotes the peptide’s approximate rounded net charge). From these, four peptides were prioritized for synthesis and *in vitro* evaluation (PIAM11-4, PIAM21-7, PIAM23-4 and PIAM29-7) hereafter referred to as PIAM11, PIAM21, PIAM23 and PIAM29, respectively. Their physicochemical properties are summarized in Table 3.

**Table 3.**
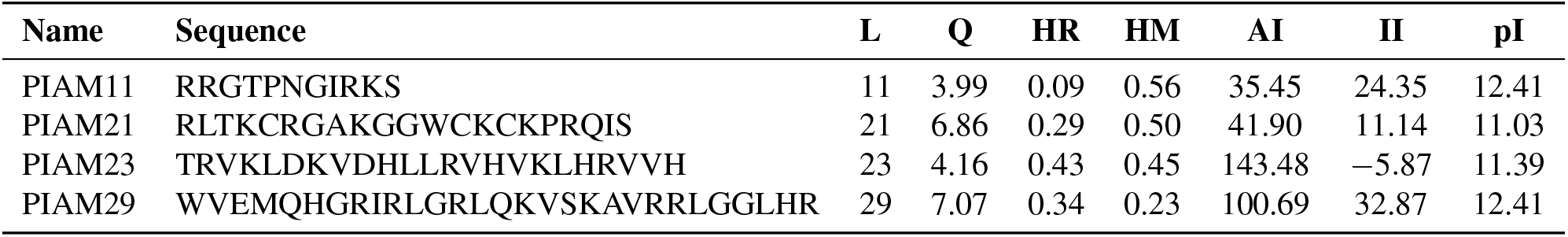
Physicochemical properties of the four selected peptide candidates. L, length (aa); Q, net charge; HR, hydrophobic ratio; HM, hydrophobic moment; AI, aliphatic index; II, instability index; pI, isoelectric point.

| Name | Sequence | L | Q | HR | HM | AI | II | pI |
| --- | --- | --- | --- | --- | --- | --- | --- | --- |
| PIAM11 | RRGTPNGIRKS | 11 | 3.99 | 0.09 | 0.56 | 35.45 | 24.35 | 12.41 |
| PIAM21 | RLTKCRGAKGGWCKCKPRQIS | 21 | 6.86 | 0.29 | 0.50 | 41.90 | 11.14 | 11.03 |
| PIAM23 | TRVKLDKVDHLLRVHVKLHRVVH | 23 | 4.16 | 0.43 | 0.45 | 143.48 | -5.87 | 11.39 |
| PIAM29 | WVEMQHGRIRLGRLQKVS KAVRRLGGLHR | 29 | 7.07 | 0.34 | 0.23 | 100.69 | 32.87 | 12.41 |

Structural predictions using PEP-FOLD3, generated under solution conditions approximating physiological pH and ionic strength, indicated that all four selected peptides predominantly adopt disordered, random-coil conformations, without a stable, well-defined secondary structure (Figure 2a–d). This contrasts with an initial round of predictions using AlphaFold, which suggested well-defined amphipathic helices for PIAM23 and PIAM29 (Supplementary Figure S2); we consider the PEP-FOLD3 predictions, obtained under explicit solvent and ionic-strength conditions closer to those used in our functional assays, more representative of the peptides’ likely conformational behavior in solution. This structural convergence was corroborated experimentally by FT-IR secondary-structure analysis of the amide I band, which likewise indicated a predominantly disordered, random-coil-dominant conformation for all four peptides in solution, with no evidence of a stable, pre-organized helix for any of them. Given the absence of a defined secondary structure across the panel, the differential antibacterial activity described below cannot be attributed to a helix-versus-coil structural dichotomy. The peptide synthesis was performed by GenScript (GenScript, USA). Purification was performed by HPLC (*>*95%) on a Vydac C18 (2.2 cm *×* 25 cm) and Luna C18 (4.6 *×* 50 mm) columns.

**Figure 2.**
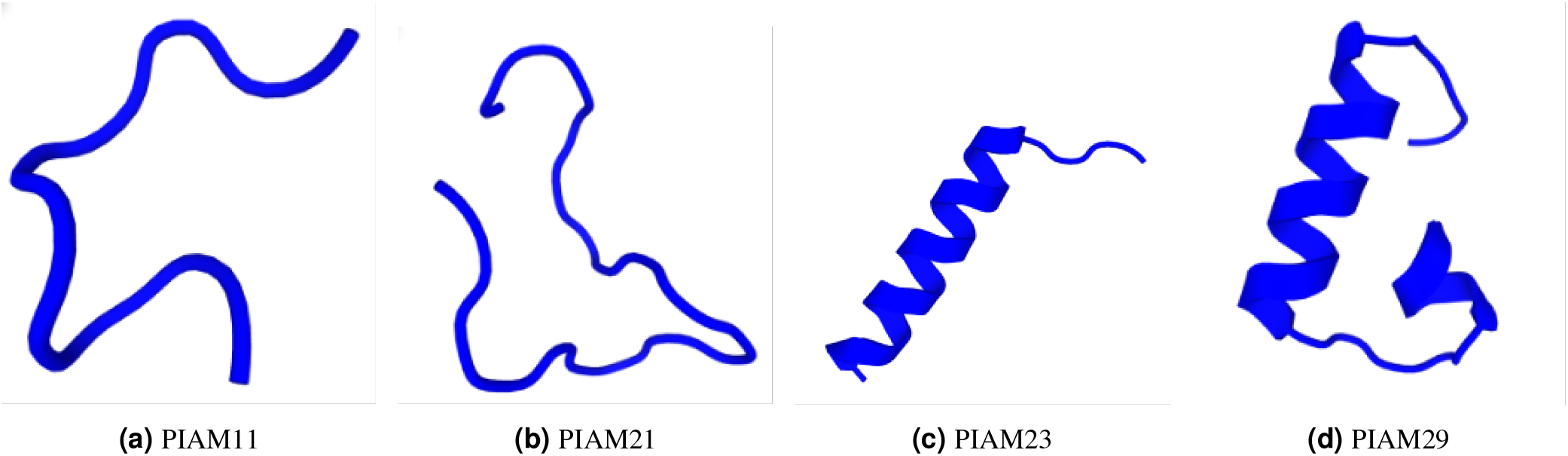
PEP-FOLD3-predicted three-dimensional structures of the four peptide candidates, generated under solution conditions approximating physiological pH and ionic strength. All four peptides are predicted to adopt predominantly disordered, random-coil conformations, without a stable, well-defined secondary structure.

### Experimental Evaluation of *M. furfur* -derived Candidate Peptides

#### Minimum Inhibitory Concentration against S. aureus

To determine whether these physicochemical differences translated into functional antimicrobial behavior, we performed kinetic broth microdilution assays following CLSI M07 guidelines^21^. These assays provided high-resolution growth curves over 24 hours, allowing us to distinguish between bacteriostatic and bactericidal effects and to evaluate concentration-dependent behavior. The results revealed that two of the four candidates presented antimicrobial activity. PIAM11 and PIAM21 showed no measurable or very poor activity across the entire concentration range (250–0.98 *µ*g/mL). Their growth curves (Figure 3a–b) overlapped with the untreated control at all timepoints, confirming the absence of bacterial growth inhibition. This behavior is consistent with numerous reports demonstrating that active AMPs require a minimum combination of cationicity and hydrophobicity to insert into bacterial lipid membranes; peptides that lack this balance typically fail to perturb membrane integrity^22^.

**Figure 3.**
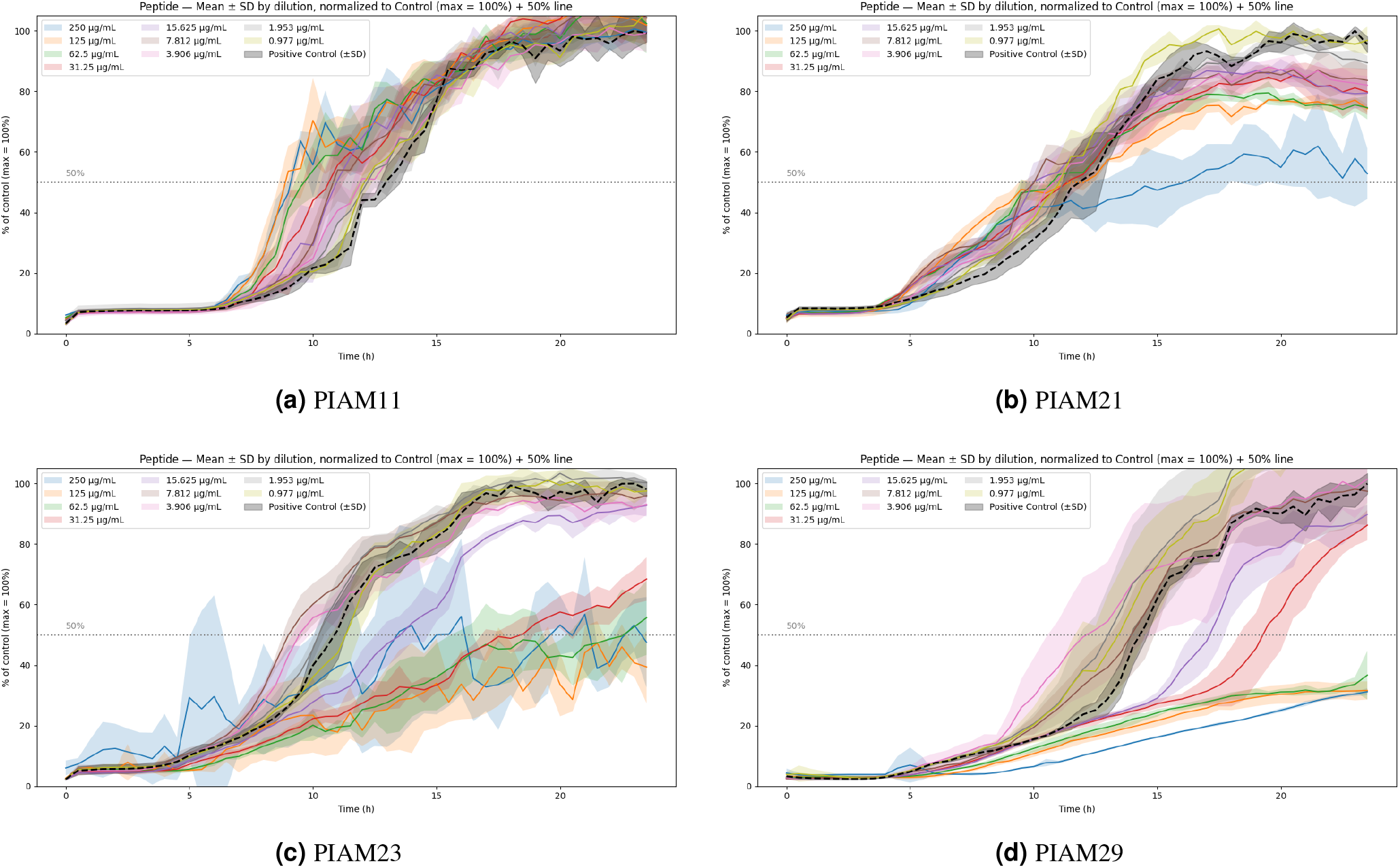
Growth curves of *S. aureus* treated with the four candidate peptides over 24 h. **(a–b)** Neither PIAM11 nor PIAM21 inhibited bacterial growth within the tested concentration range. **(c–d)** PIAM23 and PIAM29 exhibit dose-dependent bacteriostatic inhibition of *S. aureus*, with *∼*50% inhibition at 31.25 *µ*g/mL.

Conversely, PIAM23 and PIAM29 exhibited clear concentration-dependent bacteriostatic activity (Figure 3c–d). Both peptides achieved approximately 50% inhibition (MIC_50_, defined as the lowest peptide concentration maintaining growth below 50% of the growth controls over the 24-hour incubation period) at 31.25 *µ*g/mL. Their OD_600_ profiles displayed classical signatures of reversible membrane perturbation: delayed exponential growth, partial suppression, and recovery at sub-threshold concentrations^22^. Post-incubation plating confirmed that inhibited cultures remained viable, indicating a bacteriostatic rather than bactericidal effect. These findings indicate a clear differential antibacterial activity within the candidate set that, as discussed below, cannot be attributed to differences in secondary structure and is instead examined in relation to the peptides’ underlying physicochemical properties^23^.

#### Ultrastructural Profiling of S. aureus Morphology

To visually evaluate the morphological impact of the peptides on the bacterial envelope, ultrastructural profiling of *S. aureus* was performed using scanning electron microscopy (SEM) (Figure 4). Untreated control cells exhibited typical, healthy coccoid morphologies characterized by smooth, spherical, and unperturbed cell surfaces with well-defined division septa. Crucially, cells treated with the inactive candidates, PIAM11 and PIAM21, retained entirely intact and smooth cell walls, behaving indistinguishably from the untreated control.

**Figure 4.**
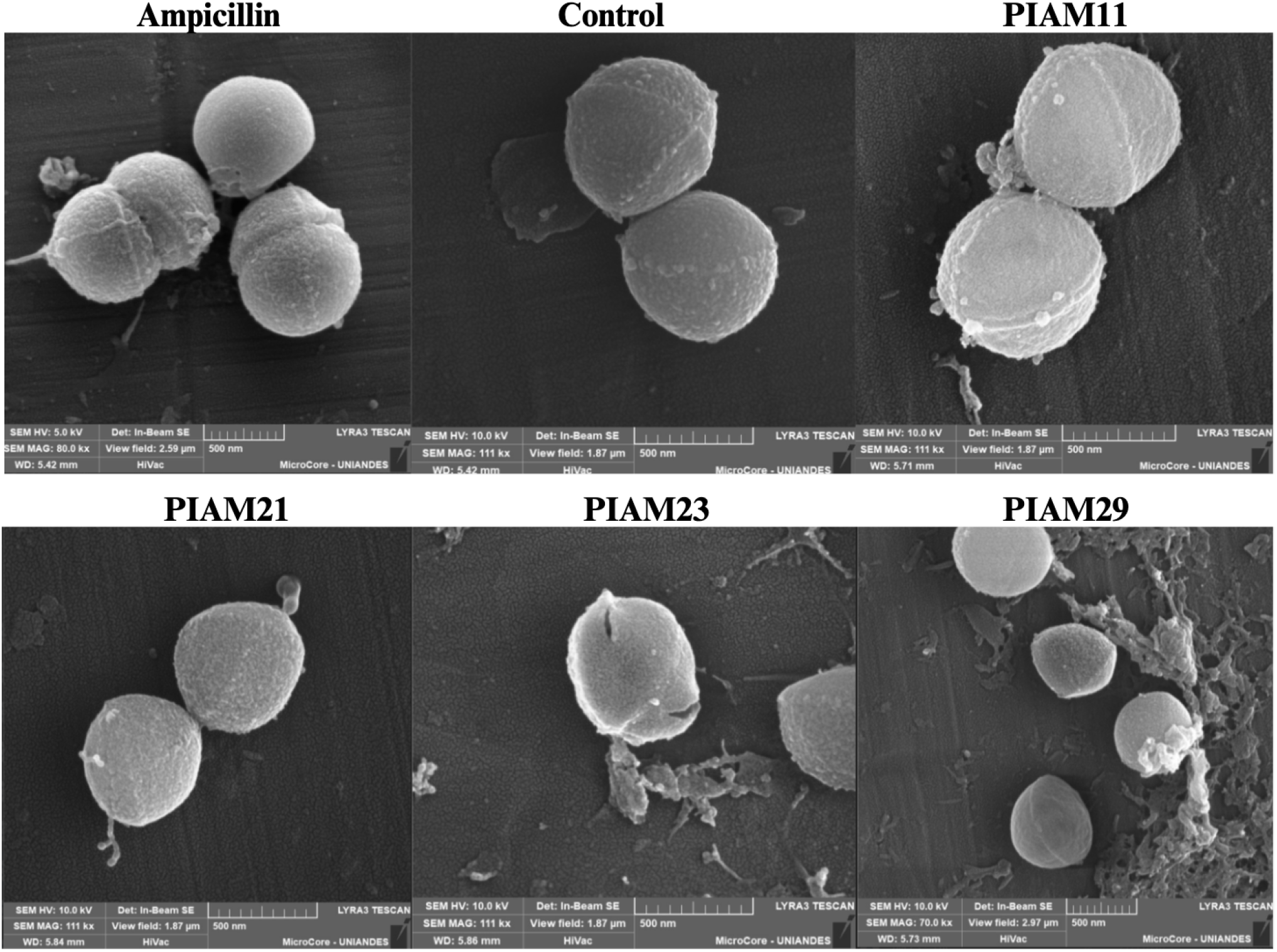
SEM micrographs of *S. aureus* (ATCC 25923). Samples treated with the active peptides (PIAM23, PIAM29) were exposed at their respective MIC_50_; samples treated with the inactive peptides (PIAM11, PIAM21), which did not reach a defined MIC_50_, were exposed at the maximum concentration used in the MIC assays (250 *µ*g/mL). The control group was left untreated, while the ampicillin group was treated with its respective MIC.

In contrast, exposure to the active peptides, PIAM23 and PIAM29, induced severe and distinct cell envelope damage. Cells treated with PIAM23 exhibited dramatic structural fracturing, characterized by deep, open clefts along the cell wall and the release of intracellular contents. Meanwhile, treatment with PIAM29 led to profound surface wrinkling, flattening, and shriveling of the cocci, accompanied by the accumulation of abundant extracellular debris consistent with envelope disruption. For comparison, ampicillin-treated cells displayed characteristic cell division defects and surface irregularities typical of cell wall synthesis inhibition, but lacked the direct, physical membrane-disruptive damage observed with the active peptides. Collectively, these ultrastructural observations are consistent with peptide-associated disruption of the *S. aureus* cell envelope and support a membrane-associated mechanism for the observed bacteriostatic activity.

#### Minimum Inhibitory Concentration against Malassezia furfur

To assess the antimicrobial spectrum and target-specificity of the designed peptides, *in vitro* susceptibility assays were conducted using the broth microdilution method in accordance with CLSI M27-A3 guidelines, with modifications to fulfill the specific nutritional requirements of *M. furfur*^24,25^. All peptides, with the exception of PIAM11, displayed detectable inhibitory activity within the tested concentration range (0.195–200 *µ*g/mL).

As shown in Table 4, PIAM29 and PIAM23 displayed the highest antifungal potency, both yielding MICs of 12.5 *µ*g/mL. PIAM21 showed moderate activity (MICs of 50–100 *µ*g/mL), while PIAM11 remained inactive (*>*200 *µ*g/mL). Notably, PIAM21 was entirely inactive against *S. aureus* (Results above) yet showed measurable antifungal activity here; because AMPs-Net prioritizes candidates from a broad, model-learned sequence space rather than from a hypothesis of activity against a specific organism, individual candidates may plausibly show target-organism-specific activity profiles that are not shared across all tested species. We consider the divergent PIAM21 profile an illustrative example of this possibility rather than an inconsistency to be resolved, and note that probing the extent to which activity profiles generalize (or fail to generalize) across target organisms is itself one of the questions this study set out to address.

**Table 4.** *In vitro* minimum inhibitory concentration (*µ*g/mL) of the candidate peptides against *M. furfur* CBS 1878.

| Peptide | MIC ($\mu\text{g/mL}$) |
| --- | --- |
| PIAM11 | >200 |
| PIAM21 | 50–100 |
| PIAM23 | 12.5 |
| PIAM29 | 12.5 |

Reference antimycotics validated the experimental model: amphotericin B and fluconazole exhibited MIC values of approximately 2 *µ*g/mL and 8 *µ*g/mL, respectively, confirming the typical susceptibility profile of *M. furfur*^25^. Overall statistical differences in antifungal potency were observed among the evaluated peptide sequences (Kruskal–Wallis, *p* = 0.0357); however, direct pairwise comparisons (Dunn’s *post-hoc* test) did not resolve statistically significant differences between individual treatments, a limitation likely driven by the small biological sample size (*n* = 3).

#### Assessment of Peptide-Induced Membrane Disruption in Lipid Bilayer Models

To further probe whether the antibacterial phenotypes observed in the kinetic broth microdilution assays were consistent with membrane-associated mechanisms, we evaluated the interaction of the peptides with a model anionic membrane using Fourier-transform infrared spectroscopy (FT-IR). We monitored the symmetric methylene stretching band (*v*_*s*_CH_2_) of DMPG bilayers as a function of temperature, which serves as a highly sensitive reporter of acyl-chain order and the gel-to-liquid crystalline phase transition^26^. In the absence of peptides (PIAM11, PIAM21, PIAM23 and PIAM29), DMPG displayed a characteristic sigmoidal thermotropic profile with a main transition centered at *∼*23.5 °C (Table 5), which aligns closely with previously reported values for pure DMPG supported lipid bilayers (SLBs)^27^.

**Table 5.** Effect of the PIAM peptides on the main transition temperatures (*T*_*m*_, °C) of DMPG SLBs. *T*_*m*_ was obtained from the *v*_*s*_CH_2_ thermotropic profiles by sigmoidal (Boltzmann) fitting. Values are mean *±* SD (SD *≈ ±*0.01 °C).

| Peptide | Control | 2.5 mol% | 5 mol% | 10 mol% |
| --- | --- | --- | --- | --- |
| PIAM11 | 23.67 | 23.72 | 23.65 | 21.11 |
| PIAM21 | 23.67 | 20.71 | 21.12 | 21.35 |
| PIAM23 | 23.67 | 21.09 | 22.24 | 22.09 |
| PIAM29 | 23.67 | 24.08 | 22.56 | 21.03 |

Upon peptide addition, all four peptides produced measurable perturbations of the DMPG thermotropic transition(Table 5), with effects that varied across peptide concentrations and were most pronounced at the highest loading of 10 mol% (Figure 5). However, the magnitude and concentration dependence of these effects did not consistently distinguish the antibacterial peptides from the inactive candidates. PIAM29 and PIAM23 (active) decreased *T*_*m*_ progressively, reaching −2.64 °C and −1.58 °C, respectively, at 10 mol%. Notably, PIAM21 (inactive) produced the largest single *T*_*m*_ depression in the entire dataset at 2.5 mol% (−2.96 °C), and PIAM11 (inactive) showed a comparable effect to the active peptides at 10 mol% (−2.56 °C) despite negligible perturbation at lower loadings.

**Figure 5.**
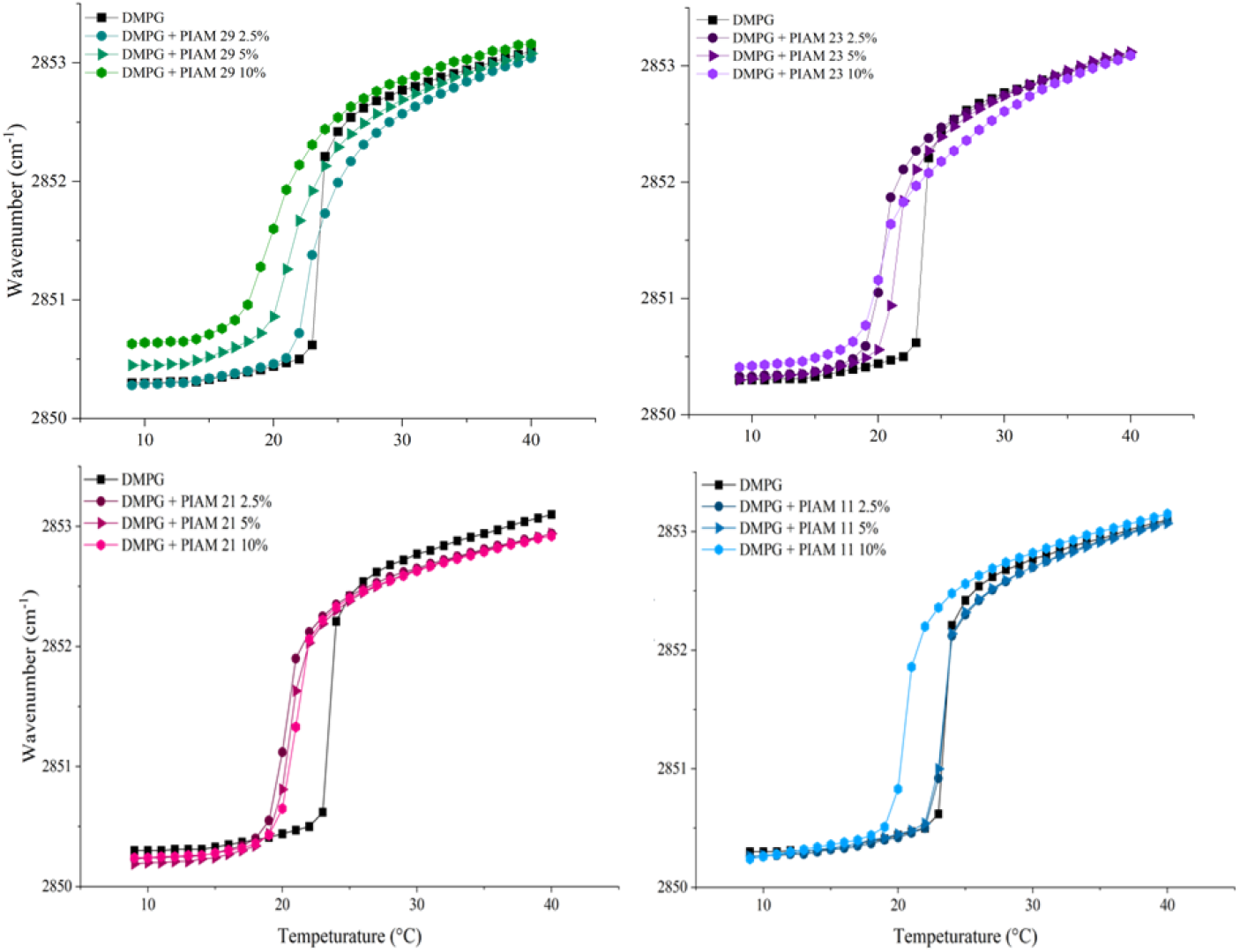
FT-IR thermotropic profiles of DMPG bilayers in the presence of PIAM peptides. Peak positions of the symmetric methylene stretching band (*v*_*s*_CH_2_) as a function of temperature for DMPG membranes with increasing peptide concentrations (2.5–10 mol%). The control exhibits a sigmoidal transition centered at *∼*23.5 °C. Active peptides (PIAM29 and PIAM23) broaden the transition and reduce cooperativity at higher loading, consistent with peptide-induced perturbation of acyl-chain packing.

These results indicate that engagement with the anionic DMPG bilayer, as measured by *T*_*m*_ depression, does not by itself predict antibacterial outcome in this candidate set: all four peptides measurably perturb the model membrane to some degree, yet only two (PIAM23 and PIAM29) produce antibacterial activity against intact *S. aureus* cells. Given that none of the four peptides adopt a stable secondary structure in solution (see above), this rules out a simple helicity-dependent mechanism and instead suggests that lipid-bilayer engagement is necessary but not sufficient for antibacterial activity in this panel, with additional factors — potentially including differences in hydrophobic content (Table 3) or behavior in the more complex environment of an intact bacterial cell — determining whether membrane interaction translates into measurable growth inhibition. We note that transition cooperativity is described here qualitatively from the shape of the fitted curves rather than through a quantitative parameter (e.g., transition width or van’t Hoff enthalpy); a formal statistical comparison against the DMPG control was likewise not performed.

#### In Vitro Cytocompatibility and Selectivity in Human Keratinocytes

To evaluate the safety profile of the identified candidate peptides (PIAM11, PIAM21, PIAM23 and PIAM29), MTT viability assays were conducted on L929 mouse fibroblasts and HaCaT human keratinocytes across a concentration range of 0.195 to 200 *µ*g/mL. Cell viability remained above 70% for all tested peptides and concentrations, satisfying the non-cytotoxicity threshold defined by ISO 10993-5:2009 (Figure 6, Supplementary Fig. S1). At the effective antimicrobial concentration (31.25 *µ*g/mL for *S. aureus* and 12.5 *µ*g/mL for *M. furfur*), cells treated with the active candidates PIAM23 and PIAM29 exhibited high viability. Similarly, exposure to PIAM11 and PIAM21 resulted in no significant reduction in cell viability across all evaluated concentrations.

**Figure 6.**
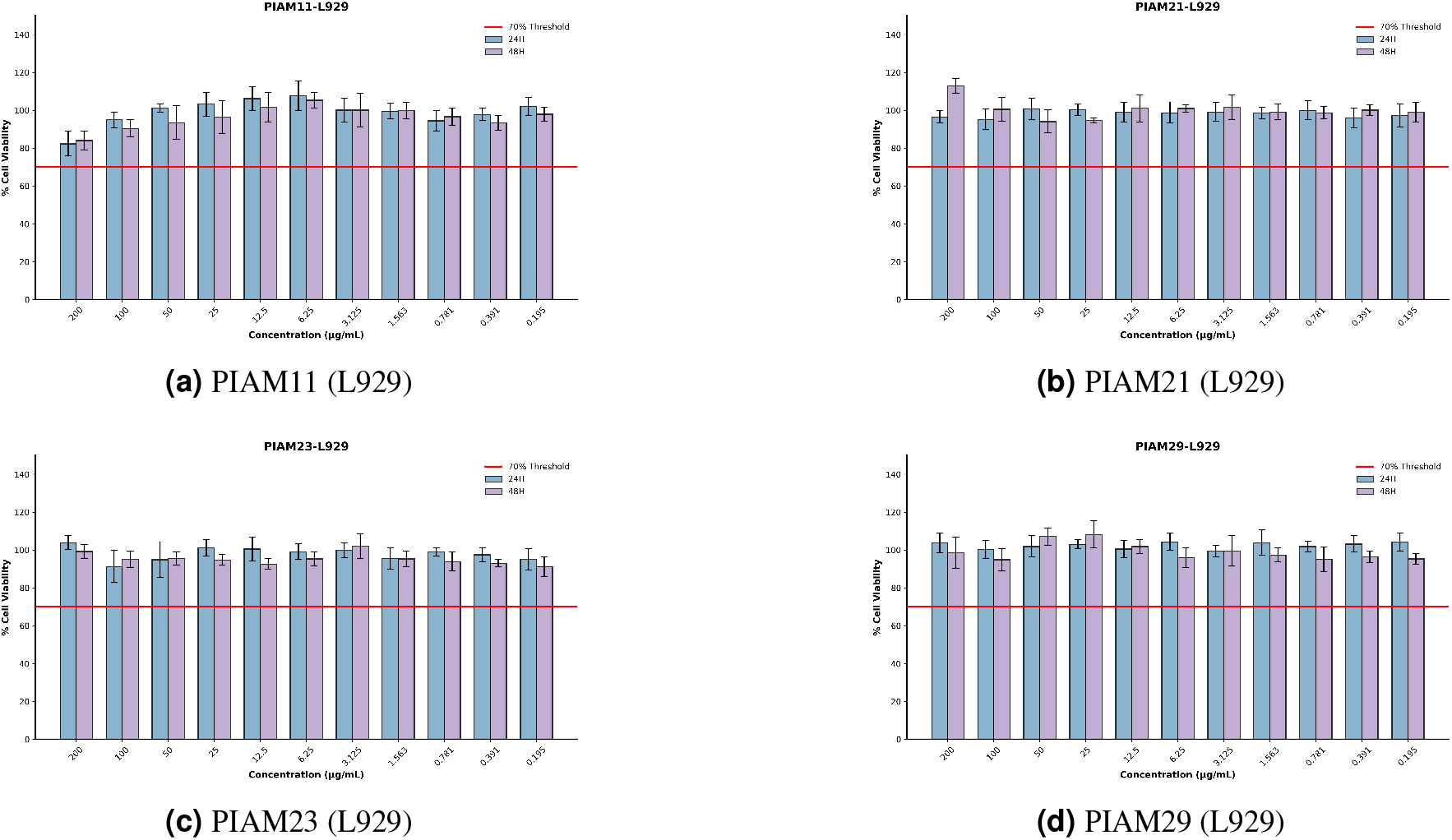
*In vitro* cytocompatibility evaluation of the four PIAM candidates in L929 mouse fibroblasts using the MTT assay. Cell viability profiles were determined after 24 h (blue bars) and 48 h (purple bars) of exposure across a concentration range from 0.195 to 200 *µ*g/mL. Panels **(a)–(d)** show PIAM11, PIAM21, PIAM23, and PIAM29, respectively. The horizontal red line denotes the 70% cell viability threshold defined by ISO 10993-5:2009 to distinguish non-cytotoxic concentrations. Data are expressed as mean *±* standard deviation.

## Discussion

This study addresses a central challenge in the translation of AI-based antimicrobial discovery: determining whether genome-guided sequence generation, combined with deep-learning prioritization, yields peptide candidates with genuine biological activity in a new, biologically motivated context^14,28^. Rather than sampling randomly from sequence space, our pipeline used the *M. furfur* genome as a constrained and ecologically justified starting point^10^. The rationale rests on an evolutionary hypothesis: that an organism sharing a resource-limited niche with *S. aureus* may have been subject to selective pressure that favors encoding antimicrobial-relevant sequence features^12,13^. This hypothesis is not one of direct secretion or functional expression of the mined fragments; on the contrary, the fragments generated through *in silico* restriction digestion do not necessarily correspond to expressed gene products. Rather, they represent a genomically bounded, biologically grounded sequence space from which AI-based inference can be applied, offering a principled alternative to unconstrained or randomly generated peptide libraries^29^. The key question this workflow poses is therefore not whether *M. furfur* naturally produces these peptides, but whether genome-anchored candidate generation enriches for sequences that AI models and subsequent experimental validation can identify as functionally active.

The computational pipeline converted an initially intractable fragment space into a tractable experimental set through successive triage steps. *In silico* restriction digestion of two *M. furfur* genomes produced 665,096 initial fragments; successive physicochemical filtering, AMPs-Net prioritization, cross-strain conservation, and antibacterial subclass selection reduced this to 37 high-confidence candidates, from which four peptides were advanced for synthesis. This funnel-like reduction provides an operational template for how AI can serve as a high-throughput prior, translating ecological hypotheses into falsifiable experimental tests^28^. Crucially, the conservation step across two strains strengthens the biological grounding of the shortlist: fragments shared across genomic backgrounds are less likely to reflect strain-specific artifacts and more likely to represent sequence determinants with structural or compositional relevance to antimicrobial function. This approach parallels recent advances in mining the human proteome for encrypted peptide antibiotics^29^, with the added layer of deep-learning-based prioritization to further refine candidate selection from a complex microbial genome.

A central concern when deploying AI-based AMP predictors in novel genomic contexts is the degree to which model performance reflects genuine biological generalizability versus dataset-specific correlations. Most publicly available AMP training datasets overrepresent particular taxonomic groups, assay conditions, and peptide families, which can introduce biases that inflate cross-validation metrics without necessarily improving prediction in out-of-distribution settings^30^. The design of our pipeline addresses this concern experimentally rather than purely computationally: by applying AMPs-Net to genome-derived fragments from an organism not represented in canonical AMP training sets, and then subjecting top-ranked candidates to multi-assay functional validation, we provide an experimental test of whether model-prioritized sequences retain measurable antimicrobial activity outside the model’s primary training distribution. The fact that two of the four synthesized candidates (PIAM23 and PIAM29) exhibited measurable antibacterial activity suggests that the model captures some generalizable features of antimicrobial potential, even when operating outside its primary training distribution. However, this observation must be interpreted cautiously. The selected peptides passed physicochemical triage and a conservation filter that already impose AMP-compatible constraints; whether AMPs-Net adds predictive value beyond these physicochemical priors, and whether its antibacterial subclass scores carry mechanistic meaning, remain open questions that motivate the structural and biophysical evidence presented below.

PEP-FOLD3 structural predictions, generated under solution conditions approximating physiological pH and ionic strength, indicated that none of the four candidates adopt a stable, well-defined secondary structure; all four are predicted to be predominantly disordered in solution. This was corroborated experimentally by FT-IR secondary-structure analysis of the amide

I band, which likewise indicated a random-coil-dominant conformation for all four peptides. An initial round of predictions using AlphaFold had suggested well-defined amphipathic helices for PIAM23 and PIAM29 (Supplementary Figure S2); given the convergence of PEP-FOLD3 (under explicit solvent and ionic-strength conditions) and the experimental FT-IR evidence, we consider the disordered conformation the more representative description of these peptides’ behavior in solution, and we no longer interpret the AlphaFold predictions as reflecting a genuine helix-versus-coil structural dichotomy within this candidate set.

Because none of the four peptides is pre-organized as a stable helix in solution, the differential antibacterial activity observed between PIAM23/PIAM29 (active) and PIAM11/PIAM21 (inactive) cannot be explained by a structural dichotomy, and must instead be examined in relation to the peptides’ underlying physicochemical properties (Table 3). Notably, the two active peptides have a substantially higher hydrophobic ratio and aliphatic index (PIAM23: HR 0.43, AI 143.48; PIAM29: HR 0.34, AI 100.69) than the two inactive peptides (PIAM21: HR 0.29, AI 41.90; PIAM11: HR 0.09, AI 35.45), with PIAM11’s hydrophobic ratio in particular markedly below that of the other three candidates. This raises the possibility that a minimum threshold of overall hydrophobic content, rather than a pre-organized amphipathic structure, is the primary physicochemical determinant of antibacterial activity within this panel. Consistent with this reframing, many membrane-active peptides remain largely disordered in bulk solvent and are thought to undergo conformational ordering only upon contact with an anionic lipid interface^31^; the disordered state observed here in solution does not preclude local, membrane-induced folding at the point of contact. This remains a hypothesis rather than a directly measured phenomenon: definitive resolution of peptide conformation at the membrane interface would require structural methods such as X-ray crystallography, which are beyond the scope of the present computational-to-experimental pipeline.

Complementing these findings, ultrastructural analysis by SEM revealed pronounced morphological alterations *S. aureus* cells following treatment with the active peptides, consistent with cell-envelope disruption. Cells treated with the inactive candidates (PIAM11 and PIAM21) remained comparable to the untreated control, maintaining smooth, unperturbed spherical surfaces. Exposure to PIAM23 and PIAM29 at their respective MIC_50_ values produced severe surface alterations, including deep cell wall wrinkling, irregular cell contours, and localized wall perforations, features indicative of severe peptide-induced envelope stress and mechanical cell wall damage^31,32^. These SEM observations are consistent with the biophysical evidence obtained via FT-IR (below), suggesting a mechanistic pathway for the active candidates in which membrane engagement triggers lipid bilayer perturbation, which ultimately manifests as structural damage at the cellular level^22,23^; as above, this pathway does not depend on a pre-organized helical conformation.

FT-IR measurements in anionic DMPG model membranes indicate that membrane engagement, unlike secondary structure, does not cleanly separate active from inactive peptides: all four peptides produced measurable, concentration-dependent perturbations of the lipid thermotropic transition, including cases where an inactive peptide (PIAM21) produced the single largest *T*_*m*_ depression in the dataset. This indicates that lipid-bilayer engagement, as captured by this assay, is necessary but not sufficient to explain antibacterial outcome, and that the distinction between active and inactive candidates in this panel likely depends on additional factors not resolved by DMPG *T*_*m*_ alone — potentially including differences in hydrophobic content (Table 3) or peptide behavior in the more complex environment of an intact bacterial cell^33^.

This pattern parallels the mode of action reported for membrane-targeting encrypted peptides from the human proteome, many of which are themselves intrinsically disordered in solution^29^, and the concentration-dependent lipid perturbation observed here is broadly consistent with the thermotropic behavior reported for other membrane-active agents in anionic bilayers^27^. Together, the structural predictions and FT-IR data suggest that neither AlphaFold-predicted helicity nor DMPG *T*_*m*_ depression alone is a sufficient predictor of antibacterial activity within this candidate set, and that hydrophobic content (Table 3) is a more parsimonious correlate of the outcomes observed here.

Extending this membrane-engagement mechanism from bacterial to fungal systems, the potent activity of PIAM23 and PIAM29 against *M. furfur* (MIC = 12.5 *µ*g/mL) highlights a compelling cross-reactivity. Rather than suggesting an endogenous secretory role in the source organism, this shared potency reflects two intersecting biophysical factors. First, both *S. aureus* and *M. furfur* present anionic membrane surfaces susceptible to electrostatic engagement by these cationic candidates. Second, fungal plasma membranes are uniquely enriched in sterols, complex lipids, and specific phospholipids (such as phosphatidylserine and phosphatidylinositol) that provide an ordered hydrophobic environment, which can significantly amplify peptide insertion relative to bacterial systems^34–36^. Conversely, surface components in Gram-positive walls (e.g., lipoteichoic acids) may partially sequester the peptides, explaining the slightly lower baseline potency observed against *S. aureus*^36^.

Ultimately, this dual antibacterial and antifungal activity underscores that hydrophobic, cationic peptides prioritized by AI from a genomic search space can engage and disrupt microbial membranes without requiring a pre-organized secondary structure^37^. While this demonstrates the power of AI to identify sequences with genuine antimicrobial potential, it also highlights a current limitation in discriminating narrow-from broad-spectrum candidates, and in disentangling the contribution of AI-based prioritization from that of the underlying physicochemical filters (see above), marking both target specificity and model attribution as key open challenges for future computational design.

The cytocompatibility profile of the active candidates (PIAM23 and PIAM29) provides encouraging signals for early-stage preclinical validation. MTT assays in L929 mouse fibroblasts and HaCaT human keratinocytes demonstrated cell viabilities well above the ISO 10993-5:2009 non-cytotoxicity threshold (*>*70%) across all evaluated concentrations (0.195–200 *µ*g/mL). Crucially, a significant quantitative separation exists between the therapeutic doses required for cross-kingdom antimicrobial activity (MIC50 = 12.5–31.25 *µ*g/mL against both *S. aureus* and *M. furfur*) and the onset of mammalian cytotoxicity, which remained undetected even at the maximum testing limit of 200 *µ*g/mL. This clear divergence establishes a favorable *in vitro* selectivity index (*SI >* 6.4), providing a foundational benchmark for preclinical safety^38^. This selective pattern is fundamentally driven by the differential lipid composition of microbial versus mammalian membranes^22,37^. While bacterial and fungal envelopes present highly anionic surfaces susceptible to electrostatic binding, mammalian bilayers contain cholesterol and are predominantly zwitterionic, offering a less permissive environment for the hydrophobic insertion of cationic peptides^22^. The absence of cytotoxicity for PIAM11 and PIAM21 is consistent with this compositional selectivity: although both peptides measurably perturb the anionic DMPG bilayer (FT-IR results above), the zwitterionic, cholesterol-rich composition of mammalian membranes appears to limit their engagement with host cell membranes regardless of overall antibacterial potency, reinforcing that membrane composition, rather than peptide activity *per se*, is the principal determinant of the observed selectivity. Collectively, these data validate a promising *in vitro* therapeutic window for PIAM23 and PIAM29, although physiologically relevant factors such as salt sensitivity, serum protein binding, and localized pH dynamics remain to be systematically evaluated in subsequent preclinical phases.

We present a reproducible computational-to-experimental strategy for discovering antimicrobial peptides encoded in the *M. furfur* genome and for evaluating the real-world validity of AI-based AMP inference. Two deep-learning–prioritized peptides exhibited dose-dependent bacteriostatic activity against *S. aureus*, perturbed anionic lipid bilayers in FT-IR assays, and induced severe mechanical wall fracturing as confirmed by SEM analysis, while remaining fully cytocompatible in mammalian skin cell lines, despite lacking a stable, pre-organized secondary structure in solution. Notably, these peptides also demonstrated potent antifungal activity against *M. furfur*, providing additional *in vitro* validation of their biological activity and supporting the broader functional relevance of the predicted sequences within their ecological origin. Together, these results support the feasibility of AI-guided genome mining as a practical route to discovery and provide a comprehensive validation framework for translating peptide inference into experimentally confirmed, selective antimicrobial candidates.

## Methods

### Computational Pipeline for Genome-Derived Peptide Discovery

#### Malassezia furfur Genome Sources

We used two publicly available *Malassezia furfur* genome assemblies as independent sources for peptide mining: strain CBS 1878 and strain 4DS^16,17^. Using two strains enabled a conservation-based filter, prioritizing peptide fragments shared across genomes and reducing the likelihood of selecting strain-specific sequences.

#### In Silico Restriction Digestion and Peptide Fragment Generation

To generate candidate fragments, both genomes were cleaved *in silico* using the restriction enzymes *CfoI* and *Sau3AI*^14,17^. We identified all genomic cut positions matching each enzyme recognition motif and extracted the resulting DNA fragments between consecutive cut sites. DNA fragments were translated into peptide sequences using the standard genetic code. To avoid fragments unlikely to be biologically meaningful as peptides, only translated sequences without internal stop codons were retained for downstream processing. Fragment sets were generated independently per strain and enzyme.

#### Physicochemical Descriptor Computation

For each peptide fragment, we computed physicochemical properties commonly used to characterize AMPs^14,19^: length (L), net charge (Q), hydrophobicity (H), hydrophobic moment (HM), instability index (II), and isoelectric point (pI). Hydrophobicity and hydrophobic moment were computed using a standard amino-acid hydrophobicity scale, and the instability index and pI were calculated using established sequence-based formulations. These descriptors were used both to filter the initial fragment pool and to guide the final downselection for synthesis.

#### Physicochemical Filtering

To enrich for fragments with AMP-like biophysical profiles, we applied a first-pass physicochemical filter prior to deep-learning inference. Briefly, we retained sequences within an AMP-compatible length window (10–50 amino acids), required overall cationicity (positive net charge), and constrained hydrophobicity-related properties to avoid highly hydrophobic or excessively amphipathic sequences that are frequently associated with poor solubility or aggregation^19,20^. The resulting candidate counts per strain and enzyme are reported in Tables 1 and 2.

#### Deep-Learning Prioritization Using AMPs-Net (Binary Classification)

Fragments passing physicochemical filtering were evaluated using the binary AMPs-Net classifier^14^, a deep-learning predictor trained to classify peptides as antimicrobial or non-antimicrobial. We retained peptides with predicted antimicrobial probability greater than 0.90. This threshold was selected to prioritize high-confidence candidates for downstream filtering while maintaining a manageable set size.

#### Cross-Strain Conservation Filtering by Exact Sequence Identity

To prioritize candidates likely to reflect conserved genomic determinants, we next retained only fragments present as an identical peptide sequence in both strains (CBS 1878 and 4DS, within each restriction-enzyme set); this is reported as “Cross-strain conservation (exact match)” in Tables 1 and 2.

#### Deep-Learning Prioritization Using AMPs-Net (Multiclass Classification)

Conserved candidates were then evaluated using the multiclass AMPs-Net classifier, which estimates subclass probabilities (antibacterial, antiviral, antifungal, and antiparasitic). We retained peptides with predicted antibacterial probability greater than 0.95, yielding the final high-confidence candidate sets summarized in Tables 1 and 2.

#### Downselection of Candidates for Synthesis

From the high-confidence antibacterial candidates, we selected a subset for synthesis based on physicochemical tractability and expected stability in aqueous handling conditions. Selection criteria considered net charge, hydrophobic moment, instability index, and peptide length^20,39^. This process produced a shortlist of seven candidates, from which four peptides (PIAM29, PIAM23, PIAM21, and PIAM11) were selected for synthesis and *in vitro* validation. The physicochemical properties of these four peptides are reported in Table 3.

#### Structure Prediction and Peptide Synthesis

To obtain structural hypotheses for the synthesized candidates, we predicted peptide three-dimensional structures using PEP-FOLD3^40^, run under solution conditions approximating physiological pH and ionic strength, matching the conditions used in our functional assays (Figure 2). For comparison, we additionally generated predictions using AlphaFold^41^, which does not explicitly model solvent pH or ionic strength (Supplementary Figure S2). Because short peptides may remain conformationally flexible in solution, all predictions were interpreted as structural tendencies rather than definitive conformations. As an experimental complement to these predictions, peptide secondary structure in solution was also assessed by FT-IR analysis of the amide I band. The synthetic peptides (PIAM11, PIAM21, PIAM23, and PIAM29; GenScript) were synthesized via solid-phase methods (purity *>*95% by analytical HPLC), with TFA removal and molecular weights confirmed by MALDI-TOF mass spectrometry.

### Minimum Inhibitory Concentration against *Staphylococcus aureus*

Antibacterial activity of PIAM11, PIAM21, PIAM23 and PIAM29 were assessed according to the Clinical and Laboratory Standards Institute guidelines M07^21^ as depicted in Figure 7a.

**Figure 7.**
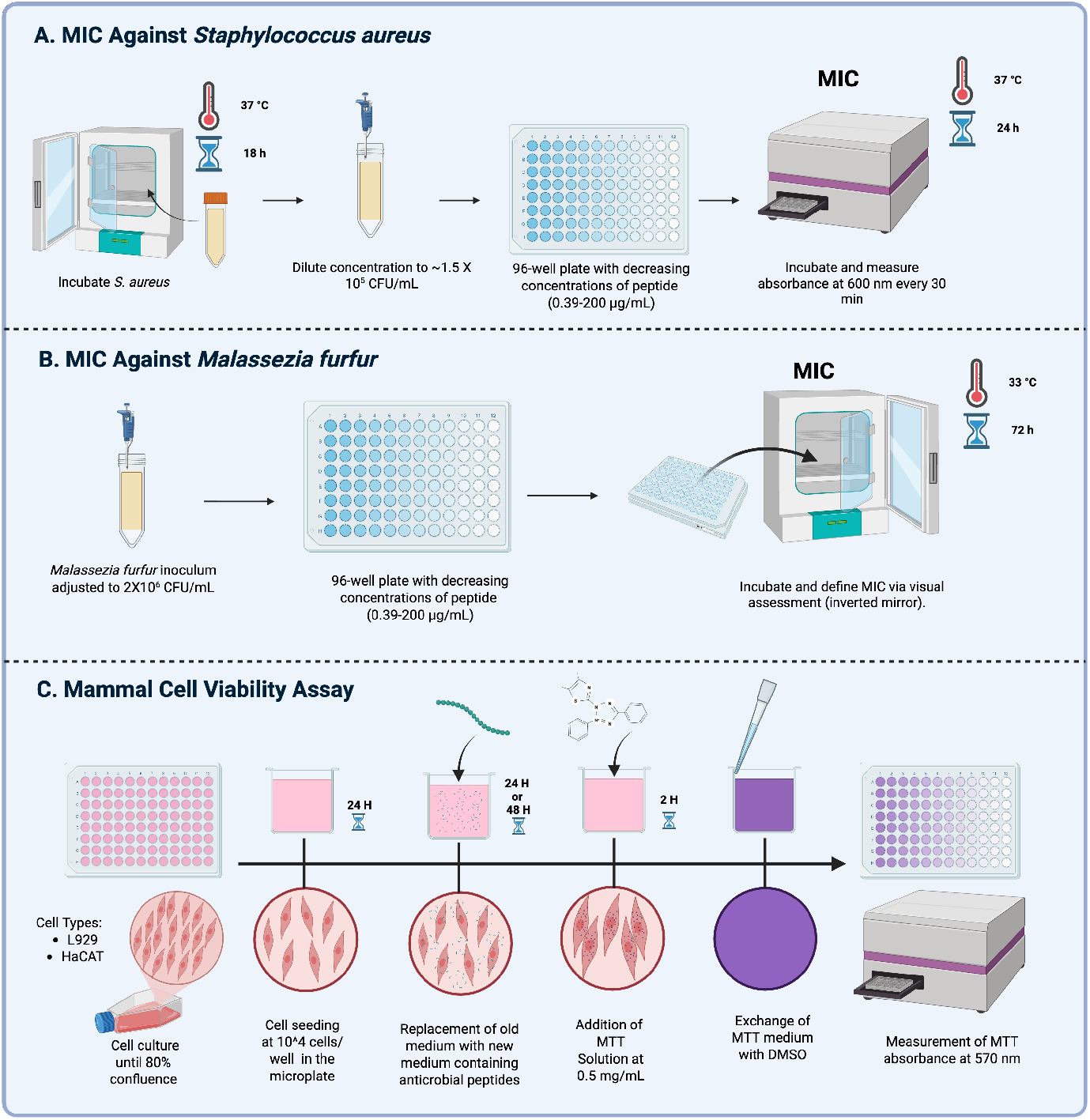
Experimental validation workflow of selected AMPs. This schematic outlines the comprehensive *in vitro* testing pipeline used to evaluate the biological efficacy and safety of the prioritized candidates across a concentration range of 0.195 to 200 *µ*g/mL. The integrated workflow comprises three core steps: **(a)** determination of the MIC against the Gram-positive bacterium *Staphylococcus aureus* via continuous kinetic absorbance monitoring at 600 nm over 24 hours; **(b)** evaluation of targeted antifungal activity by establishing the MIC against *Malassezia furfur* via visual assessment following a 72-hour incubation period; and **(c)** assessment of mammalian cytocompatibility in L929 mouse fibroblasts and HaCaT human keratinocytes using the standard MTT colorimetric assay (0.5 mg/mL) at 24 and 48 hours post-exposure.

Initially, we screened AMP inhibitory activity against *S. aureus* (ATCC 25923). To determine antibacterial activity and MIC50 values, the peptides were tested over a nine-step two-fold dilution series in Type II water, spanning from a maximum concentration of 250 *µ*g/mL to a minimum of 0.98 *µ*g/mL.

Bacteria were inoculated in Mueller–Hinton broth and incubated at 37 °C overnight to reach log phase. The cell suspension was diluted to achieve a final inoculum density in accordance with CLSI M07 requirements (*∼*5 *×* 10^5^ CFU/mL). Then, 50 *µ*L aliquots were transferred into 96-well plates containing 50 *µ*L of peptide solution.

Peptide solutions at different concentrations were monitored using a multimodal microplate reader operating at 37 °C for 24 h, with optical density at 600 nm (OD_600_) measurements recorded every 30 min. Growth curves were plotted, and antibacterial activities of the peptides were assessed.

MIC values were determined as the lowest peptide concentration maintaining growth comparable to sterility controls throughout the full 24-hour incubation period. MIC_50_ values were defined as the lowest peptide concentration that maintained growth below 50% of the growth controls over the 24-hour incubation period.

### Scanning Electron Microscopy

SEM was employed to characterize *S. aureus* membrane morphology following previous methodology^32^. Bacterial suspensions were adjusted to an optical density of 0.2 (OD_600_) and incubated for 2 h at 37 °C with peptide solutions at their respective MIC_50_ (PIAM23, PIAM29) or, for the peptides that did not reach a defined MIC_50_ (PIAM11, PIAM21), at the maximum concentration used in the MIC assays (250 *µ*g/mL), as well as with ampicillin at its MIC. Subsequently, 10 *µ*L of samples were deposited onto a coin covered with sterile aluminum foil and allowed to air dry. The samples were then analyzed at the Microscopy Center (MicroCore, *µ*-core), Office of the Vice President for Research and Creation, Universidad de los Andes, by SEM observation using a Tescan LYRA 3 scanning electron microscope (TESCAN, Brno, Czech Republic).

### Minimum Inhibitory Concentration against *Malassezia furfur*

Antifungal susceptibility assay was performed following CLSI M27-A3 guidelines as depicted in Figure 7b, with minor adaptations to the protocol to assess *M. furfur* nutritional requirements^24,25,42^. Briefly, the four AMP candidates PIAM11, PIAM21, PIAM23 and PIAM29 were tested in a 96-well round-bottom microplate at concentrations ranging from 0.195 to 200 *µ*g/mL. Each peptide was initially dissolved in sterile distilled water to obtain a stock concentration of 2000 *µ*g/mL and subsequently diluted in Sabouraud dextrose broth supplemented with 0.5% Tween 60. Appropriate growth and sterility controls were included in all experiments. Concentrations ranging from 0.03 to 16 *µ*g/mL of fluconazole (FCZ) and amphotericin B (AMB) were used as internal controls to verify the susceptibility and reproducibility of the *M. furfur* strain to antifungal compounds.

The *M. furfur* inoculum was prepared in sterile distilled water containing 0.5% Tween 80 and diluted in accordance with CLSI M27-A3 requirements to a final inoculum density of 1 *×* 10^3^ CFU/mL per well, using a Neubauer counting chamber to verify inoculum density. Plates were incubated at 33 °C for 72 hours and visually assessed using an inverted mirror. The MIC was defined as the lowest peptide concentration resulting in *≥*90% growth inhibition compared to the untreated control.

### Fourier-Transform Infrared Spectroscopy (FT-IR) of Peptide–Membrane Interactions

To probe peptide-induced perturbations on a Gram-positive membrane-mimetic system, we monitored the thermotropic phase behavior of supported lipid bilayers (SLBs) composed of 1,2-dimyristoyl-*sn*-glycero-3-phosphoglycerol sodium salt (DMPG; Avanti Polar Lipids) as was already done in previous work^27^. DMPG was selected because C14 acyl chains are reported among the most abundant in *Staphylococcus aureus* membranes. To form the SLBs, 20 *µ*L of a 20 mM DMPG stock solution in chloroform was deposited onto the silicon crystal of a BioATR II cell. Following solvent evaporation, the lipid film was hydrated with 20 *µ*L of buffer (20 mM HEPES, 500 mM NaCl, 1 mM EDTA; pH 7.4) at 37 °C for 10 min. Peptides were introduced during this hydration step at final concentrations corresponding to 2.5–10 mol% relative to the lipid.

Lipid phase-transition measurements were performed using a Tensor II FT-IR spectrometer (Bruker Optics) equipped with a mercury cadmium telluride (MCT) detector. Temperature was precisely controlled (accuracy *±*0.01 °C) using a Huber Ministat 125 circulating water bath. Background spectra of the buffer were recorded during a programmed heating ramp from 7 to 40 °C at a rate of 1 °C/min, with a 120 s equilibration time between measurements. Spectral processing was conducted in OPUS 3D. After automatic background subtraction and baseline correction (20% sensitivity rubber-band method), phase transitions were tracked by monitoring the symmetric methylene stretch band (*v*_*s*_CH_2_) in the 2850–2853 cm^−^1 region. The maximum wavenumber at each temperature was extracted using the OPUS peak-picking tool, and the resulting wavenumber-versus-temperature profiles were fitted to a Boltzmann sigmoidal function using a Levenberg–Marquardt iteration to determine the main phase transition temperature (*T*_*m*_) at the inflection point. All experiments were performed in triplicate to assess repeatability.

### Viability Assessment by MTT Assay

Cell viability was evaluated using the standard MTT colorimetric assay in accordance with ISO 10993-5:2009 guidelines^43^ as depicted in Figure 7c. Briefly, L929 mouse fibroblast (ATCC) and HaCaT human keratinocyte cells were seeded in 96-well plates at a density of 1 *×* 10^4^ cells/well and incubated for 24 h under standard culture conditions to allow for adherence. The peptide candidates (PIAM11, PIAM21, PIAM23, and PIAM29) were initially dissolved in 30 mM HEPES buffer (pH 7.0–7.4) to obtain a 1 mg/mL stock solution. Working solutions were prepared immediately prior to cell treatment via serial dilution in Dulbecco’s Modified Eagle Medium (DMEM), and the cells were subsequently exposed to 11 distinct peptide concentrations ranging from 0.195 to 200 *µ*g/mL for periods of 24 h and 48 h.

Following the exposure time, the treatment medium was replaced with 100 *µ*L of fresh medium containing 0.5 mg/mL

MTT, and the plates were incubated for 4 h at 37 °C. After carefully aspirating the supernatant, the resulting insoluble purple formazan crystals were solubilized by adding 100 *µ*L of dimethyl sulfoxide (DMSO) to each well. Absorbance was then measured at 570 nm using a microplate spectrophotometer. Cell viability was calculated as a percentage by normalizing the absorbance values of the treated wells to those of the untreated control cells (established as 100% viability). All experimental conditions were performed in quadruplicate and repeated across two independent experiments.

### Statistical Analysis

All experimental biological assays, including antimicrobial kinetic screenings, antifungal susceptibility testing, and mammalian cell viability evaluations, were performed in at least two or three independent replicates, each conducted with technical triplicates to ensure reproducibility. Quantitative data are expressed as mean *±* standard deviation (SD). Statistical significance across the experimental groups and peptide sequences was evaluated using non-parametric analysis; specifically, a Kruskal–Wallis test was employed for multiple comparisons, followed by Dunn’s *post-hoc* test to resolve variance between individual treatment pairs. A *p*-value of less than 0.05 (*p <* 0.05) was considered statistically significant. All computational statistical evaluations, curve fitting, and data visualizations were performed using GraphPad Prism software (v8.0) and custom Python routines.

## Acknowledgements

This research was funded by the Colombian Ministry of Science, Technology, and Innovation (Minciencias), under the 937-2023 Call for Fundamental Research. This work was supported by Azure sponsorship credits granted by Microsoft’s AI for Good Research Lab.

## Author contributions statement

S.O. conceived the genome-mining strategy, developed the computational pipeline (*in silico* restriction digestion, physicochemical filtering, and AMPs-Net inference), and wrote the corresponding sections of the manuscript. P.Av. performed the minimum inhibitory concentration assays against *S. aureus* and contributed to the adaptation of the experimental protocol, and wrote the corresponding section. S.C. performed the MTT cytocompatibility assays and wrote the corresponding section. V.R.R. performed the FT-IR membrane-interaction experiments and wrote the corresponding section. P.L. performed the minimum inhibitory concentration assays against *M. furfur* and wrote the corresponding section. M.M.M. and C.L. supervised the FT-IR and biophysical membrane experiments. A.M.C.R. supervised the *Malassezia*-related experimental work. C.M.C., C.L. and P.Ar. conceived and supervised the study. All authors reviewed and edited the manuscript.

## Data availability

The *M. furfur* CBS 1878 and 4DS genome assemblies analyzed in this study are publicly available, as described in Triana *et al*.^16^. AMPs-Net, the deep-learning model used for antimicrobial peptide prioritization, is publicly available at https://github.com/BCV-Uniandes/AMPs-Net. All experimental data generated in this study (growth curves, MIC values, FT-IR thermotropic profiles, and MTT viability data) are available within the article and its Supplementary Information, or from the corresponding author upon request.

