## Supplementary Information for "Assessing the translation of AI-prioritized genome-derived peptide fragments into validated antimicrobial candidates"

Sebastian Ojeda[1], Paula Avila[1], Stiven Castellanos[1], Paloma Lemaitre[2], Valeria Ruiz-Ramírez[4], Marcela Manrique-Moreno[4], Adriana Marcela Celis Ramírez[2], Pablo Arbeláez[1], Chad Leidy[3], and Carolina Muñoz-Camargo[1,*]

[1]Department of Biomedical Engineering, Universidad de los Andes, Bogotá, Colombia
[2]Department of Biological Sciences, Universidad de los Andes, Bogotá, Colombia
[3]Department of Physics, Universidad de los Andes, Bogotá, Colombia
[4]Chemistry Institute, Universidad de Antioquia - UdeA, Medellín, Colombia
[*]

| Name | Sequence | L | Q | HR | HM | AI | II | pI |
|---|---|---|---|---|---|---|---|---|
| PIAM29-7 | WVEMQHGRIRLGRLQKVSKAVRRLGGLHR | 29 | 7.07 | 0.34 | 0.23 | 100.69 | 32.87 | 12.41 |
| PIAM23-3 | AASLTFFKARTLTRTVAGLASNH | 23 | 3.03 | 0.48 | 0.26 | 85.22 | 8.40 | 12.13 |
| PIAM23-4 | TRVKLDKVDHLLRVHVKLHRVVH | 23 | 4.16 | 0.43 | 0.45 | 143.48 | −5.87 | 11.39 |
| PIAM21-4 | KHGVGMCERRGNAYGNRVHRL | 21 | 3.92 | 0.29 | 0.45 | 50.95 | 11.65 | 11.11 |
| PIAM21-6 | RLTKCRGAKGGWCKCKPRQIS | 21 | 6.53 | 0.29 | 0.12 | 41.90 | 11.14 | 11.03 |
| PIAM18-5 | IGQGLVKVKCKPVGTKKG | 18 | 4.83 | 0.33 | 0.18 | 91.67 | −6.11 | 10.86 |
| PIAM11-4 | RRGTPNGIRKS | 11 | 3.99 | 0.09 | 0.56 | 35.45 | 24.35 | 12.41 |

Table 1: Amino acid sequences and physicochemical properties of the seven candidate peptides identified through the AI-guided genome-mining pipeline. L, length (aa); Q, net charge; HR, hydrophobic ratio; HM, hydrophobic moment; AI, aliphatic index; II, instability index; pI, isoelectric point. Four of these (PIAM11-4, PIAM21-4, PIAM23-4, and PIAM29-7) were selected for synthesis and experimental validation, hereafter referred to in the main text as PIAM11, PIAM21, PIAM23, and PIAM29, respectively.

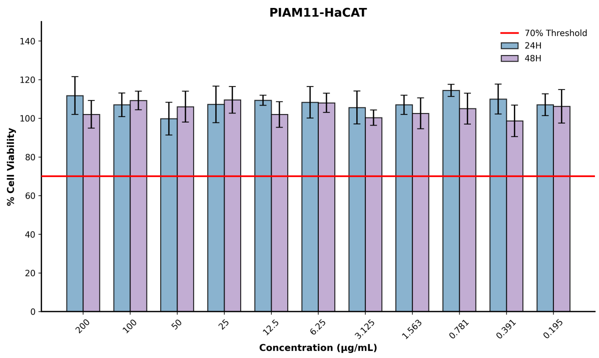


(a) PIAM11

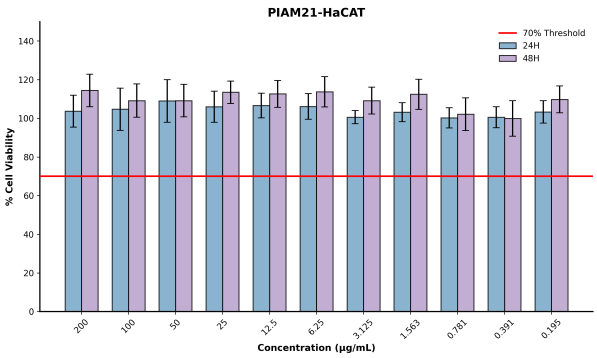


(b) PIAM21

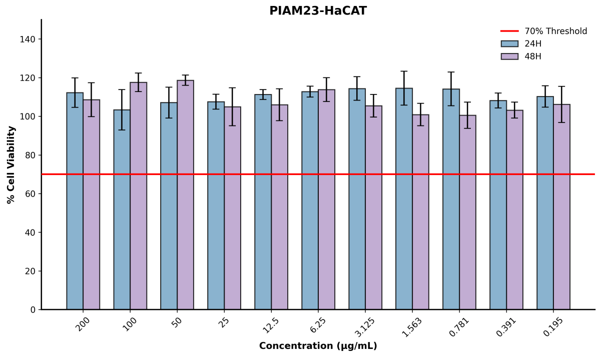


(c) PIAM23

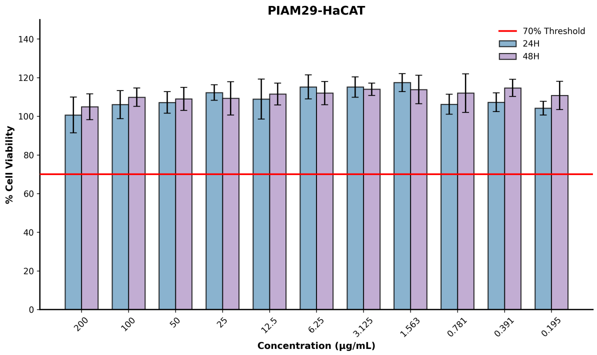


(d) PIAM29

Figure 1: Representative microscopy images of HaCaT human keratinocytes following exposure to the four PIAM candidates. Images are shown for PIAM11, PIAM21, PIAM23, and PIAM29 and provide morphological evidence complementary to the quantitative cell-viability measurements obtained using the MTT assay.

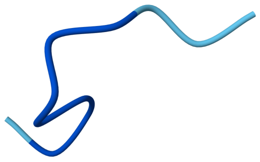

(a) PIAM11

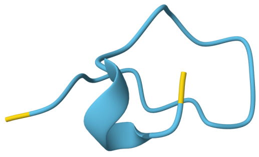

(b) PIAM21

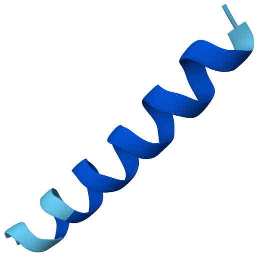

(c) PIAM23

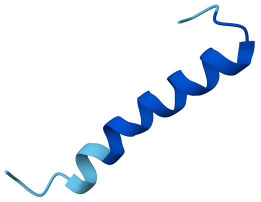

(d) PIAM29

Figure 2: AlphaFold-predicted three-dimensional structures of the four peptide candidates, shown here for comparison only. Under these predictions, PIAM23 and PIAM29 appeared as well-defined amphipathic helices, whereas PIAM11 and PIAM21 lacked defined secondary structure. However, PEP-FOLD3 predictions generated under solution conditions approximating physiological pH and ionic strength (Figure 2, main text), corroborated by FT-IR secondary-structure analysis, indicate that all four peptides predominantly adopt disordered, random-coil conformations. The AlphaFold predictions shown here are therefore not considered representative of the peptides' behavior under the conditions used in this study.
